# Perception of environmental polypeptides in *C. elegans* activates insulin/IGF signaling and alters lipid metabolism

**DOI:** 10.1101/341883

**Authors:** Rebecca E. W. Kaplan, Amy K. Webster, Rojin Chitrakar, Joseph A. Dent, L. Ryan Baugh

**Author notes:** Corresponding author, L. Ryan Baugh Department of Biology, Duke University, Box 90338, Durham, NC 27708-0338.

## Abstract

Food perception affects animal physiology in complex ways. We uncoupled the effects of food perception and ingestion in the roundworm *C. elegans*. Perception was not sufficient to promote development, but larvae exposed to food without ingestion failed to develop upon return to normal culture conditions. Inhibition of gene expression during perception rescued subsequent development, demonstrating the response to perception without feeding is deleterious. Perception altered DAF-16/FOXO localization, reflecting activation of insulin/IGF signaling (IIS). The insulin-like peptide *daf-28* was specifically required, suggesting perception in chemosensory neurons directly regulates peptide secretion. Gene expression and Nile Red staining suggest that perception alters lipid metabolism. Environmental polypeptides are sensed by starved larvae and promote dauer diapause recovery. We conclude that polypeptides are perceived as a food-associated cue, initiating a signaling and gene regulatory cascade that alters metabolism in anticipation of feeding and development, but that this response is detrimental if feeding does not occur.

## Introduction

Perception of food affects metabolism and development in a variety of animals. Several observations suggest that sensory perception of food can regulate metabolism. For example, humans release insulin in response to the sight and smell of food (Sjostrom, Garellick et al. 1980). In mice loss of olfactory neurons reduces obesity and insulin resistance, and enhancing olfactory acuity does the reverse (Riera, Tsaousidou et al. 2017). Blocking olfaction in *Drosophila* alters metabolism and extends lifespan; conversely, the longevity-extending effects of dietary restriction are partially reversed by exposure to food odors (Libert, Zwiener et al. 2007). Likewise, in *C. elegans* sensory perception affects lifespan and development of dauer larvae, a form of diapause in the third larval stage (Apfeld and Kenyon 1999, Alcedo and Kenyon 2004, Hu 2007, Lans and Jansen 2007). However, the molecular cues that are sensed and their specific effects on organismal signaling and gene regulation are not well understood in any system.

*C. elegans* L1-stage larvae hatch in a state of developmental arrest (“L1 arrest” or “L1 diapause”) and require food to initiate development (Baugh 2013). IIS is a key regulator of L1 arrest (Baugh and Sternberg 2006, Fukuyama, Rougvie et al. 2006). During starvation, *daf-16*/FOXO promotes L1 arrest by inhibiting development-promoting pathways (Baugh and Sternberg 2006, Kaplan, Chen et al. 2015). Feeding up-regulates activity of the insulin-like peptides *daf-28*, *ins-6*, and *ins-4*, among others, which act as agonists for the only known insulin/IGF receptor *daf-2* (Chen and Baugh 2014). *daf-2*/InsR signaling activates a conserved phosphoinositide 3-kinase (PI3K) cascade to antagonize DAF-16 and promote development (Morris, Tissenbaum et al. 1996, Lin, Dorman et al. 1997, Ogg, Paradis et al. 1997, Kimura, Riddle et al. 2011). *daf-28*, *ins-6*, and *ins-4* are also critical to regulation of dauer development (Li, Kennedy et al. 2003, Cornils, Gloeck et al. 2011), which together with their role in regulating L1 development, indicates that they are atop the organismal regulatory network governing postembryonic development. However, how these important insulin-like peptides are regulated in response to nutrient availability is unknown.

Feeding in *C. elegans* is mediated by pumping of the neuromuscular organ called the pharynx (Avery and You 2012). The drug ivermectin paralyzes the pharynx by activating glutamate-gated chloride channels containing α-type channel subunits, increasing chloride conductance and inhibiting cellular depolarization (Avery and Horvitz 1990, Cully, Vassilatis et al. 1994, Dent, Davis et al. 1997, Vassilatis, Arena et al. 1997). Several genes encoding glutamate-gated chloride channels in *C. elegans* confer sensitivity to ivermectin, but simultaneous mutation of three or more of these genes produces substantial ivermectin resistance (Dent, Smith et al. 2000).

Here we used ivermectin to prevent feeding in worms exposed to food. We show that perception of food without ingestion significantly alters gene expression and activates IIS but is not sufficient to initiate development. To the contrary, perception without ingestion makes developmental arrest irreversible. We show that starved worms sense polypeptides in their environment as a food cue, likely in anticipation of feeding.

## Results

### Perception of food without ingestion renders developmental arrest irreversible

We used ivermectin to prevent feeding in order to uncouple the effects of food perception from ingestion. We wanted to limit effects of the drug outside the pharynx, so we started with a highly ivermectin-resistant strain, the quadruple mutant *avr-14(vu47); glc-3(ok321) avr-15(vu227) glc-1(pk54)*, and rescued *avr-15* with a *myo-2* promoter for pharynx-specific expression. We made two versions of the strain with two different markers for analysis of development: AJM-1::GFP to examine seam cells and *Phlh-8::GFP* to examine the M-cell lineage.

Throughout this study, most experiments follow the basic setup seen in Fig. 1A. We prepared embryos by hypochlorite treatment and cultured them in either ivermectin or control (DMSO) conditions without food for 24 hr so they hatch and enter L1 arrest. Various types of food or other substances were then added, and worms were typically analyzed 1 hr or 24 hr after this addition. To determine if ingestion was occurring, GFP beads were added to the cultures and worms were examined. Critically, GFP beads were not ingested in the ivermectin plus food (*E. coli* HB101) conditions (Supp. Fig. 1A). For initial characterization of the effects of food perception without ingestion, worms were plated in standard laboratory conditions (on plates with *E. coli* OP50 but no ivermectin) after 24 hr of exposure to experimental conditions and allowed to recover for three days. Worms exposed to ivermectin plus food failed to recover, remaining arrested in the L1 stage, while the controls recovered completely (Fig. 1B). That is, ivermectin treatment alone did not cause an irreversible arrest, but ivermectin plus food did. This striking phenotype was further characterized with a time series, revealing a near complete effect by about 8 hr (Fig. 1C). Recovery to the L4 stage was chosen as an easy stage to reliably score. Though 24 hr exposure generally rendered larvae capable of negligible if any growth, earlier time points associated with incomplete penetrance were associated with intermediate growth rates as well. Worms displayed significant failure to recover with as little as 1 mg/mL HB101 (Supp. Fig. 1B), and worms were at least as sensitive to *E. coli* OP50 and HT115 (Supp. Fig. 1C). Together these results reveal a potent effect of exposure to *E. coli* without feeding on the ability of larvae to recover from starvation-induced developmental arrest.

**Figure 1.**
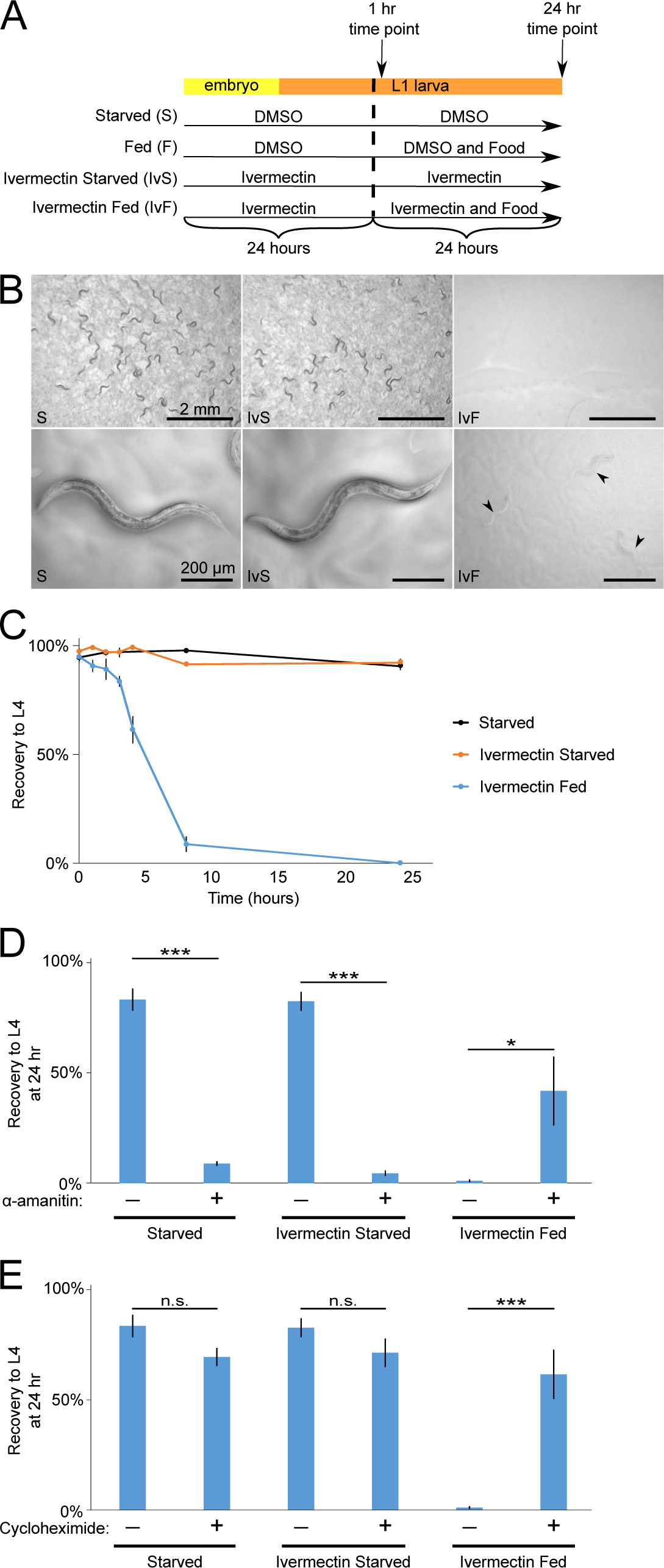
Prolonged exposure to food perception triggers an inability to recover that is mediated by a transcriptional/translational response. (A) Diagram of experimental set-up with four standard treatment conditions over time. Dimethyl sulfoxide (DMSO; solvent) (B) Representative images of worm recovery after three days post starvation. (C) L1 starvation recovery is plotted over time for three biological replicates. (D) The proportion of larvae that recovered to at least the L4 stage after three days of recovery is plotted for three to five biological replicates. (E) The proportion of larvae that recovered to at least the L4 stage after three days of recovery is plotted for three to five biological replicates. (C-E) ***p<0.001, **p<0.01; unpaired t-test. Error bars are SEM.

Feeding causes a significant change in transcription and translation in *C. elegans* L1 larvae (Baugh, Demodena et al. 2009, Maxwell, Antoshechkin et al. 2012, Stadler and Fire 2013). We hypothesized that food perception evokes a gene expression response that is deleterious without feeding. Worms were treated with the drug α-amanitin, which inhibits transcription (Sanford, Golomb et al. 1983, McColl, Rogers et al. 2010, Zaslaver, Baugh et al. 2011), slightly before and during food exposure. Blocking transcription significantly increased recovery (Fig. 1D). As a complementary approach, worms were treated with cycloheximide to block translation (McColl, Rogers et al. 2010) in a similar manner. This treatment also significantly improved recovery (Fig. 1E). Together these results suggest that food perception alters gene expression, and that this change in expression affects the animal adversely if it is not accompanied by feeding.

### Food perception evokes a gene expression response similar to feeding

We performed mRNA-seq to characterize the effects of food perception on gene expression. We assayed larvae that were exposed to ivermectin and food for 1 hr or 24 hr to distinguish relatively immediate and long-term effects, and we assayed larvae exposed to ivermectin without food at the same time points for reference, as well as larvae that were fed or starved without ivermectin for 1 hr (a 24 hr time point was not included since the fed larvae would have developed to the L3 stage). Principal component analysis revealed a large effect of ivermectin, with ivermectin treatment correlating with the first component (Supp. Fig. 2). Feeding significantly affected mRNA expression, as expected, and the second and third principal components separated the fed and starved worms (Fig. 2A). Notably, worms exposed to ivermectin and food for 1 hr were different from worms starved with ivermectin, falling closer to fed worms on the graph. However, by 24 hr of exposure to ivermectin and food the expression profile was not significantly different from its starved control. Likewise, 1,258 genes were differentially expressed at 1 hr comparing ivermectin with food to ivermectin starved, but only 241 genes were differentially expressed in the same comparison at 24 hr (false discovery rate (FDR) < 0.05 and an absolute log_2_ fold change of greater than 0.5; S1 Dataset). These results show that perception of food alters gene expression initially but that this effect subsides over time.

**Figure 2.**
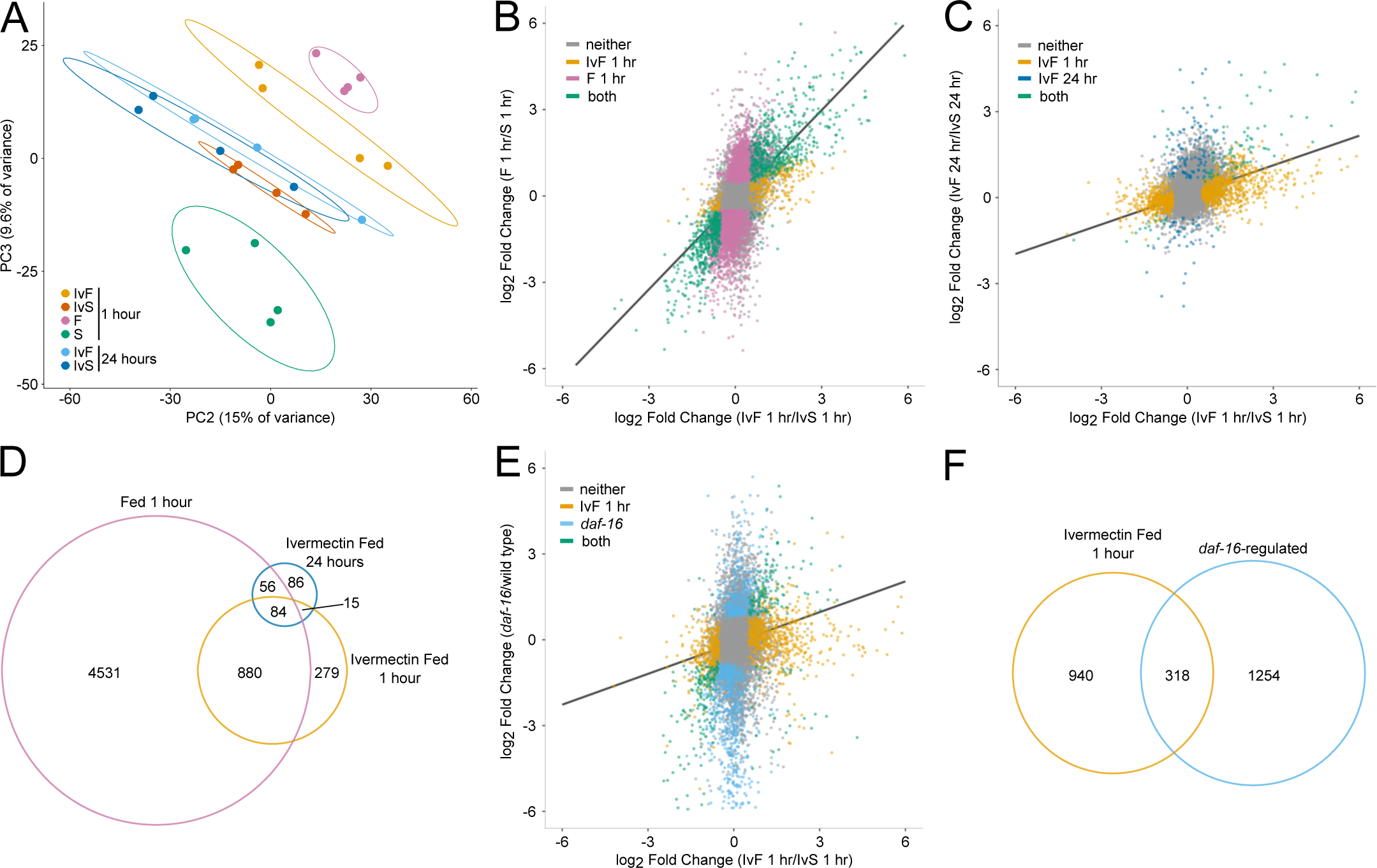
mRNA-seq reveals transcriptional effects of food perception. (A) PCA of four biological replicates is plotted. Ellipses represent 80% confidence intervals, the probability for two of which not overlapping by chance is approximately 0.04. (B) Mean gene expression changes of four biological replicates is plotted. (C) Overlap between genes significantly affected by food with genes significantly affected by ivermectin and food is plotted. (D) Mean gene expression changes of four biological replicates is plotted. (E) Overlap between genes significantly affected by ivermectin and food at 1 hr with genes significantly affected by ivermectin and food at 24 hr is plotted. (F) Mean gene expression changes of two to four biological replicates is plotted. The universal set of genes considered includes only those analyzed in both studies. (G) Overlap between genes significantly affected by ivermectin and food with genes significantly affected by daf-16 is plotted.

We wondered how well correlated the gene expression response to food perception is with feeding. The magnitude of the feeding response was larger, with 5,551 differentially expressed genes at 1 hr compared to 1,258 genes in the presence of ivermectin. These gene expression changes were very well correlated, with 98.8% of genes differentially expressed in both conditions changing in the same direction (Fig. 2B). Indeed, the vast majority of genes affected by food with ivermectin were also affected by feeding (Fig. 2D, hypergeometric p-value = 6.8e-353). These results indicate that perception of food evokes a similar, though reduced, gene expression response to feeding. The response to food in the presence of ivermectin at 1 hr and 24 hr was also well correlated, with 91.9% of genes differentially expressed at both times responding in the same direction (Fig. 2C). Indeed, there was significant overlap in the differentially expressed genes at both time points (Fig. 2D, hypergeometric p-value = 3.7e-60). These results support the conclusion that perception of food initially alters gene expression in a way that resembles the feeding response, but that that this response to perception diminishes over time.

As an effector of IIS, *daf-16*/FOXO is an important regulator of gene expression during L1 starvation (Kaplan, Chen et al. 2015, Hibshman, Doan et al. 2017). Since DAF-16 is inactivated by IIS in response to feeding, we hypothesized that it is also inactivated by perception of food, contributing to the resulting gene expression response. A previous study identified 1,572 genes differentially expressed in a *daf-16* null mutant compared to wild type during L1 starvation (Kaplan, Chen et al. 2015). These differences in gene expression correlated with the effect of food in the presence of ivermectin at 1 hr, with 88.4% of the genes significantly affected in both comparisons responding in the same direction (Fig. 2E). There was also significant overlap in the genes affected in both comparisons (Fig. 2F, hypergeometric p-value = 2.5e-98). These results suggest that perception of food in starved larvae reduces *daf-16*/FOXO activity.

### Perception of food activates insulin/IGF signaling

Similarity in the gene expression responses of wild-type worms exposed to food in the presence of ivermectin and a starved *daf-16*/FOXO mutant suggest that perception of food activates IIS. Since IIS regulates subcellular localization of DAF-16 (Henderson and Johnson 2001), perception of food should affect localization if this hypothesis is correct. We categorized GFP::DAF-16 localization as nuclear, intermediate, or cytoplasmic (Fig. 3A). As expected, GFP::DAF-16 was primarily nuclear during starvation and primarily cytoplasmic after 1 hr of exposure to food (Fig. 3B). One hour exposure to food with ivermectin also significantly shifted GFP::DAF-16 to the cytoplasm, supporting our hypothesis that perception of food activates IIS. However, after 24 hr there was no difference between ivermectin fed and ivermectin starved worms. Similar to mRNA-seq results at 1 hr and 24 hr, this result suggests that perception of food is sufficient to shift DAF-16 localization initially but not to maintain it. GFP::DAF-16 localization also responds to other bacterial strains in the presence of ivermectin (Supp. Fig. 3A). These results suggest that perception of each of the bacteria used as food in the lab can activate IIS.

**Figure 3.**
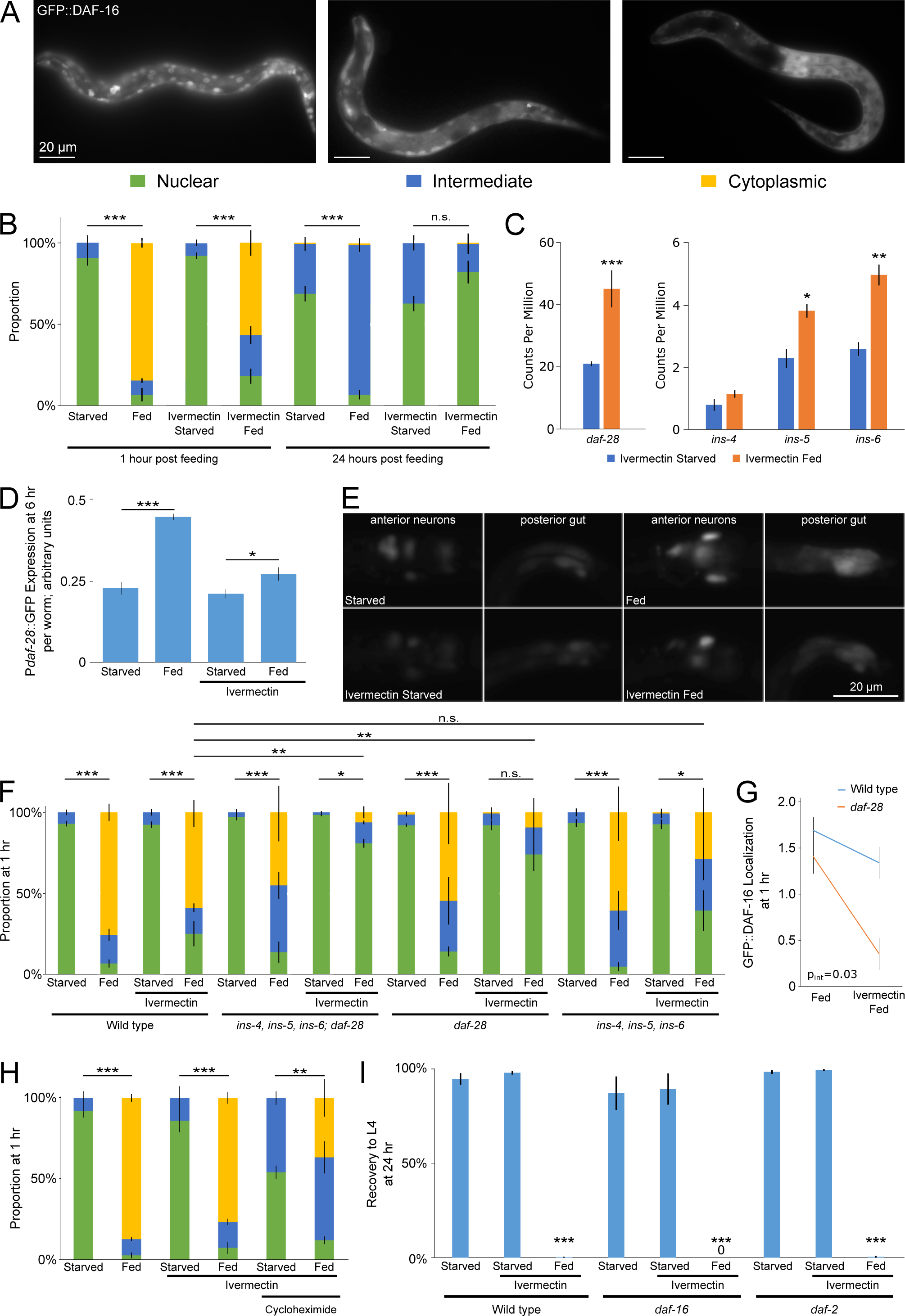
GFP::DAF-16 localization response to food perception is insulin-dependent. (A) Representative images of how GFP::DAF-16 localization was characterized. (B) GFP::DAF-16 localization is plotted for three biological replicates. (C) Average transcript abundance from four biological replicates of mRNA-Seq is plotted for selected insulin-like peptides. Nominal p-values displayed. (D) Averages of P*daf-28*::GFP fluorescence intensity normalized by optical extinction per worm using the COPAS BioSorter are plotted for four biological replicates. Exposure to HB101 was 6 hr. (E) Representative images of P*daf-28*::GFP transcriptional reporter gene are presented. (F) GFP::DAF-16 localization is plotted for three to six biological replicates. (G) GFP::DAF-16 localization is plotted for three to six biological replicates. The 2-way ANOVA interaction p-value is listed. (H) GFP::DAF-16 localization is plotted for three biological replicates. (I) The proportion of larvae that recovered to at least the L4 stage after three days of recovery is plotted for three to four biological replicates. (B-I) ***p<0.001, **p<0.01; unpaired t-test. Error bars are SEM, except for in D where they are standard deviation.

The insulin-like peptides *daf-28, ins-4, ins-5* and *ins-6* are transcriptionally up-regulated by feeding L1 larvae, and they promote L1 development (Chen and Baugh 2014). We found that *daf-28, ins-5,* and *ins-6* transcripts were significantly up-regulated after 1 hr exposure to food in the presence of ivermectin (Fig. 3C). The COPAS BioSorter was used to quantify whole-worm fluorescence of a P*daf-28*::GFP transcriptional reporter, supporting the conclusion that daf-28 transcription increases in response to food perception (Fig. 3D). This reporter was expressed in anterior neurons and the posterior intestine with brighter expression after 6 hr feeding (Fig. 3E), as expected (Chen and Baugh 2014). Consistent with the COPAS result and mRNA-seq, it was also brighter after exposure to food in the presence of ivermectin. These results reveal transcriptional up-regulation of *daf-2*/InsR agonists as an initial response to perception of food, consistent with activation of IIS.

The *C. elegans* genome encodes 40 insulin-like peptides, and many of them are functionally redundant, making it difficult to detect mutant phenotypes (Pierce, Costa et al. 2001). As a control, mutation of *daf-2*/InsR completely blocked the effects of food on GFP::DAF-16 localization (Supp. Fig. 3B). *ins-4, 5* and *6* are clustered on chromosome II, so we analyzed a deletion allele that removes all three (Hung, Wang et al. 2014), combining it with a *daf-28* deletion allele to simultaneously disrupt all four. The compound mutant retained the response to feeding, but the change in localization of GFP::DAF-16 in response to food in the presence of ivermectin was significantly reduced (Fig. 3F). A *daf-28* deletion alone mimicked the behavior of the compound mutant, but the *ins-4, 5, 6* deletion alone did not, suggesting *daf-28* specifically mediates the response to food perception. To examine this closer, we plotted the data for wild type and the *daf-28* mutant separately, focusing on the effect of food in the presence and absence of ivermectin (Fig. 3G). These data show a specific effect of *daf-28* on the response to food in the presence of ivermectin (two-way ANOVA p-values for interaction between genotype and presence or absence of ivermectin: *daf-28* = 0.03, *ins-4, 5, 6; daf-28* = 0.02, *ins-4, 5, 6* = 0.29). These data suggest that *daf-28* plays a critical role in mediating the initial response to food perception on IIS activity, and they suggest that overlapping function of insulin-like peptides provides a more robust response to feeding than perception alone.

We hypothesized that perception of food promotes secretion of DAF-28 and other insulin-like peptides from chemosensory neurons, providing a rapid response to environmental conditions. To test this hypothesis, we treated worms with cycloheximide to block translation and examined GFP::DAF-16 localization. Localization was significantly more cytoplasmic in response to food in the presence of ivermectin when treated with cycloheximide (Fig. 3H), consistent with perception of food directly promoting secretion of insulin-like peptides. However, the shift in GFP::DAF-16 localization appeared incomplete with cycloheximide treatment. Together with our results showing an effect on *daf-28* transcription, this observation suggests that food perception affects insulin-like peptide activity at multiple levels of regulation.

Given the effects of food perception on IIS, we hypothesized that IIS mutants affect the irreversible arrest resulting from perception without feeding. However, neither *daf-2*/InsR nor *daf-16*/FOXO mutants had increased recovery after exposure to food in the presence of ivermectin (Fig. 3H). If cytoplasmic localization of DAF-16 during starvation was sufficient to cause the irreversible arrest phenotype, then *daf-16* mutants should not be able to recover following starvation. *daf-16* mutants are starvation-sensitive, but they nonetheless can be starved and retain the ability to recover upon feeding (Fig. 3I). In conclusion, the irreversible arrest is likely caused by alteration of multiple pathways such that activation of IIS alone during starvation is not sufficient.

### Perception of food is not sufficient to promote development

We used Gene Ontology (GO) term enrichment analysis of our mRNA-seq results to get a broad view of the processes affected by perception of food. The response to feeding for 1 hr revealed significant overlap with metabolism genes (hypergeometric p-value = 8.6e-92) and larval development genes (Fig. 4A, hypergeometric p-value = 9.8e-39). The response to food exposure for 1 hr in the presence of ivermectin also revealed overlap with metabolism genes (Fig. 4B, hypergeometric p-value = 5.0e-23) but not with larval development genes (hypergeometric p-value = 0.89). Furthermore, genes differentially expressed in response to food exposure in the presence of ivermectin but not feeding were enriched for lipid metabolic terms, while genes differentially expressed in response to feeding but not exposure to food in the presence of ivermectin were enriched for a variety of terms related to development (Supp. Fig. 4A-C, S1 Dataset). These results suggest that perception of food affects lipid metabolism but not development.

**Figure 4.**
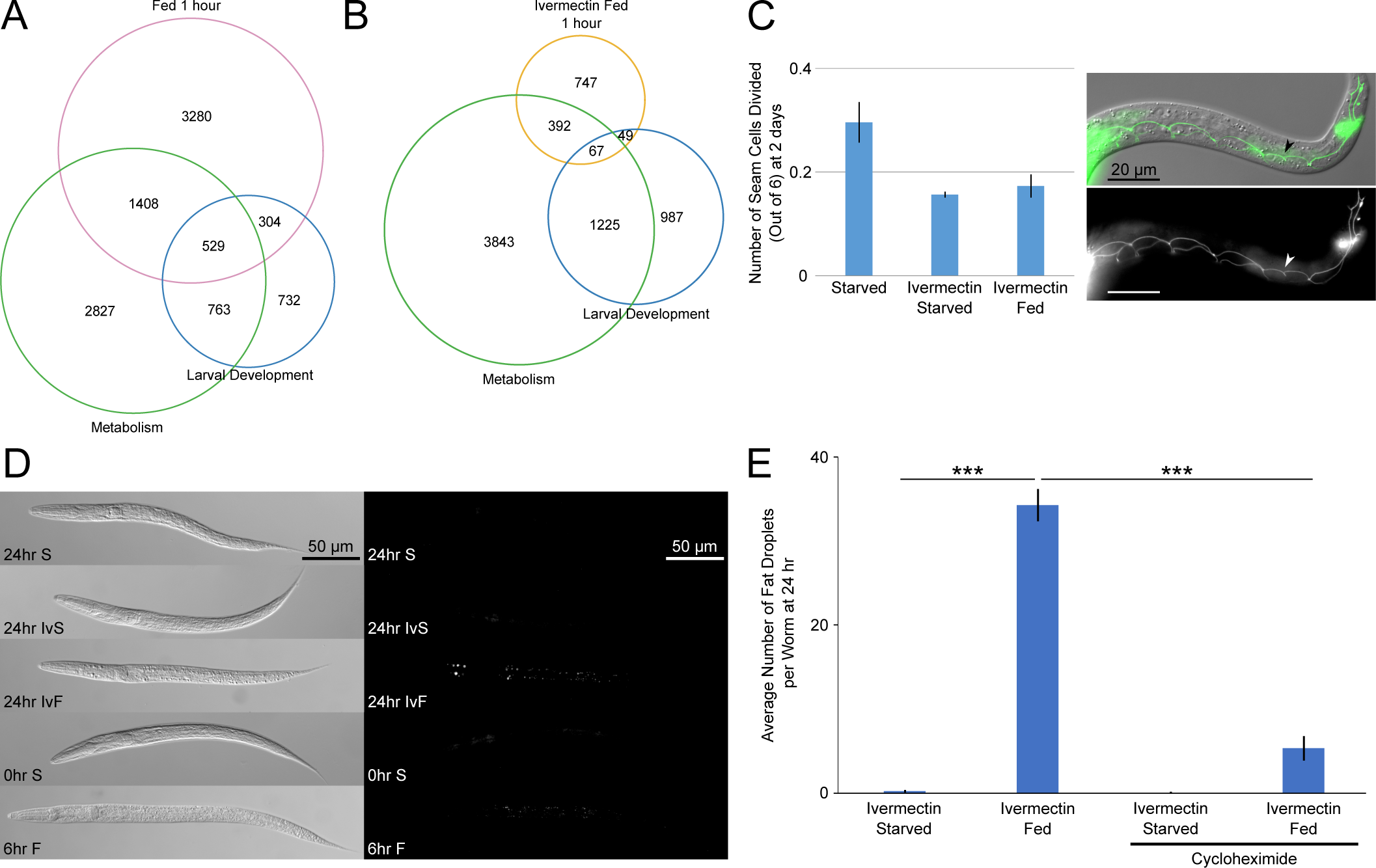
Food perception significantly affects metabolism but not development. (A) Overlap between genes significantly affected by food with genes in the metabolic process and larval developmental process GO terms is plotted. (B) Overlap between genes significantly affected by ivermectin and food with genes in the metabolic process and larval developmental process GO terms is plotted. (C) The average number of seam cell divisions, out of six possible, is plotted for three biological replicates. Scoring was done two days after HB101 addition. (D) Representative DIC and GFP channel images of fixed worms following Nile red staining are presented. (E) Quantification of fat droplets in Nile red staining is plotted for three to four biological replicates. (C-E) ***p<0.001, **p<0.01; unpaired t-test. Error bars are SEM.

The lateral epidermal seam cells are the first cells to divide in developing L1 larvae (Sulston and Horvitz 1977), and they divide very rarely during L1 arrest (Baugh and Sternberg 2006, Kaplan, Chen et al. 2015). We used an AJM-1::GFP reporter for adherens junctions to visualize seam cell membranes and count divisions of the cells v1-6 (Gupta, Wang et al. 2003). Consistent with the results of GO term analysis, exposure to food for two days in the presence of ivermectin did not cause seam cell divisions (Fig. 4C). There were also no M-cell divisions (data not shown). These results with the most stringent assay available indicate that perception of food is not sufficient to promote detectable postembryonic development.

### Perception of food alters lipid metabolism

GO term enrichments suggest that lipid metabolism is affected by perception of food. Consistent with this hypothesis, differential interference contrast microscopy revealed numerous droplets throughout the body and around the pharynx after prolonged exposure to food in the presence of ivermectin (Fig. 4D). Given their appearance and GO term enrichments (Supp. Fig. 4A), we hypothesized that these are lipid droplets. Nile red staining of fixed L1 larvae supported this hypothesis (Fig. 4D). Starved L1 larvae, either shortly after hatching or 24 hr later, did not contain such fat droplets. Fed L1 larvae developed small fat droplets in what appeared to be the intestine, while the droplets in worms exposed to food and ivermectin for 24 hr were more varied in size and location. Cycloheximide treatment significantly reduced the number of fat droplets (Fig. 4E). We conclude that the gene expression response to food perception alters lipid metabolism, resulting in abnormal accumulation of lipid droplets in the body cavity.

### Polypeptides serve as an environmental cue for food

Worms rely on mechanosensory and chemosensory cues to regulate locomotion, development, pathogen avoidance, feeding, and mating (Bargmann 2006, Goodman 2006). Worms respond to mechanosensory stimulus when encountering a bacterial lawn, which can be mimicked with Sephadex beads (Sawin, Ranganathan et al. 2000). To test whether the effects of food perception were due to mechanosensation or chemosensation, we assayed the ability to recover after starvation in the presence of ivermectin and Sephadex beads or HB101 bacterial filtrate, respectively. We found that Sephadex beads did not affect starvation recovery, while HB101 filtrate prevented recovery as strongly as HB101 itself (Fig. 5A, Supp. Fig. 1A). These data suggest that the deleterious effect of food perception without ingestion is via chemosensation and not mechanosensation.

**Figure 5.**
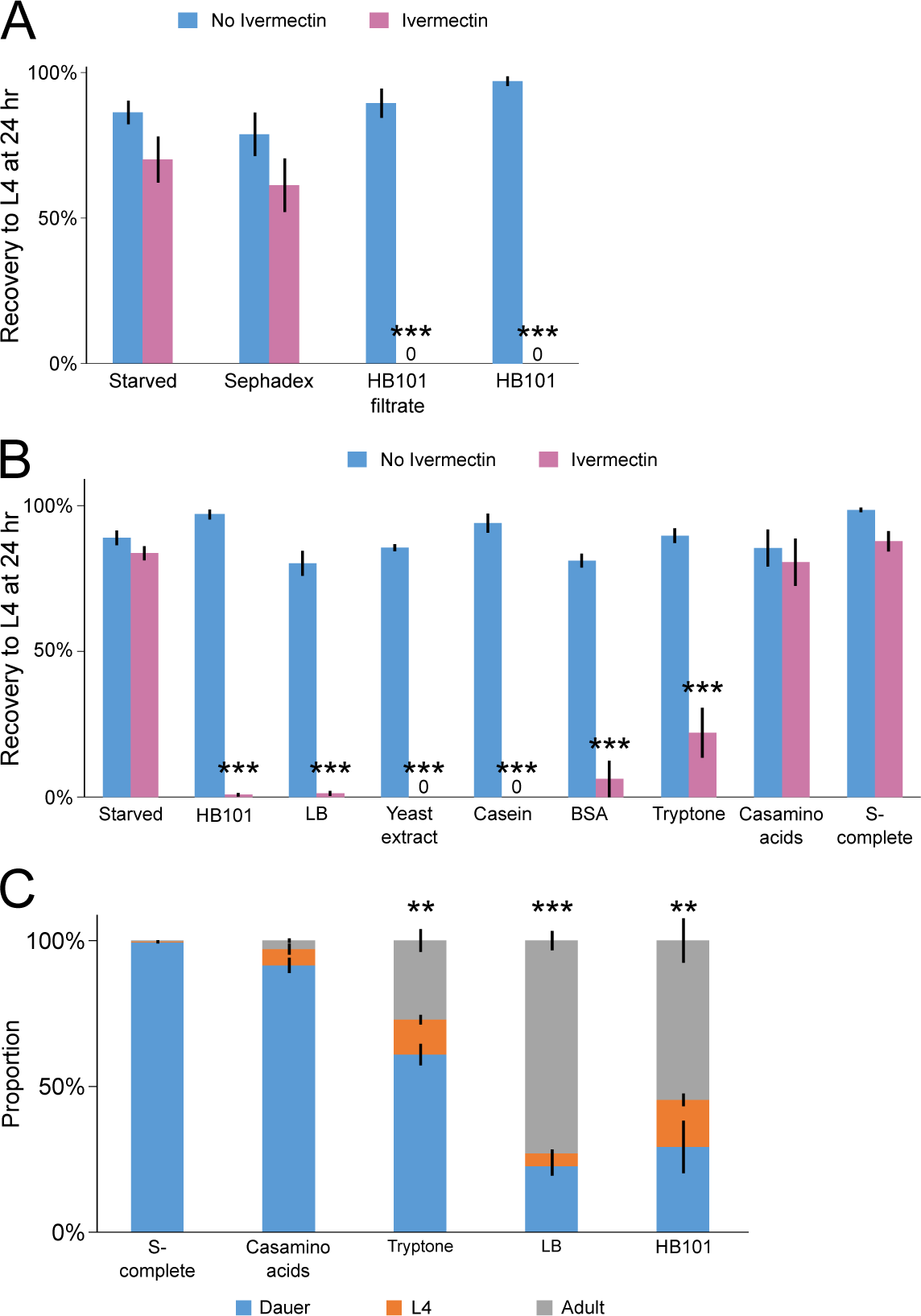
Perception of food cues affects L1 and dauer recovery. (A-B) The proportion of larvae that recovered to at least the L4 stage after three days of recovery is plotted for three to thirteen biological replicates. (C) Recovery from dauer after three days in each condition is plotted for three to four biological replicates. (A-C) ***p<0.001, **p<0.01, *p<0.05; unpaired t-test. Error bars are SEM.

**Figure 6.**
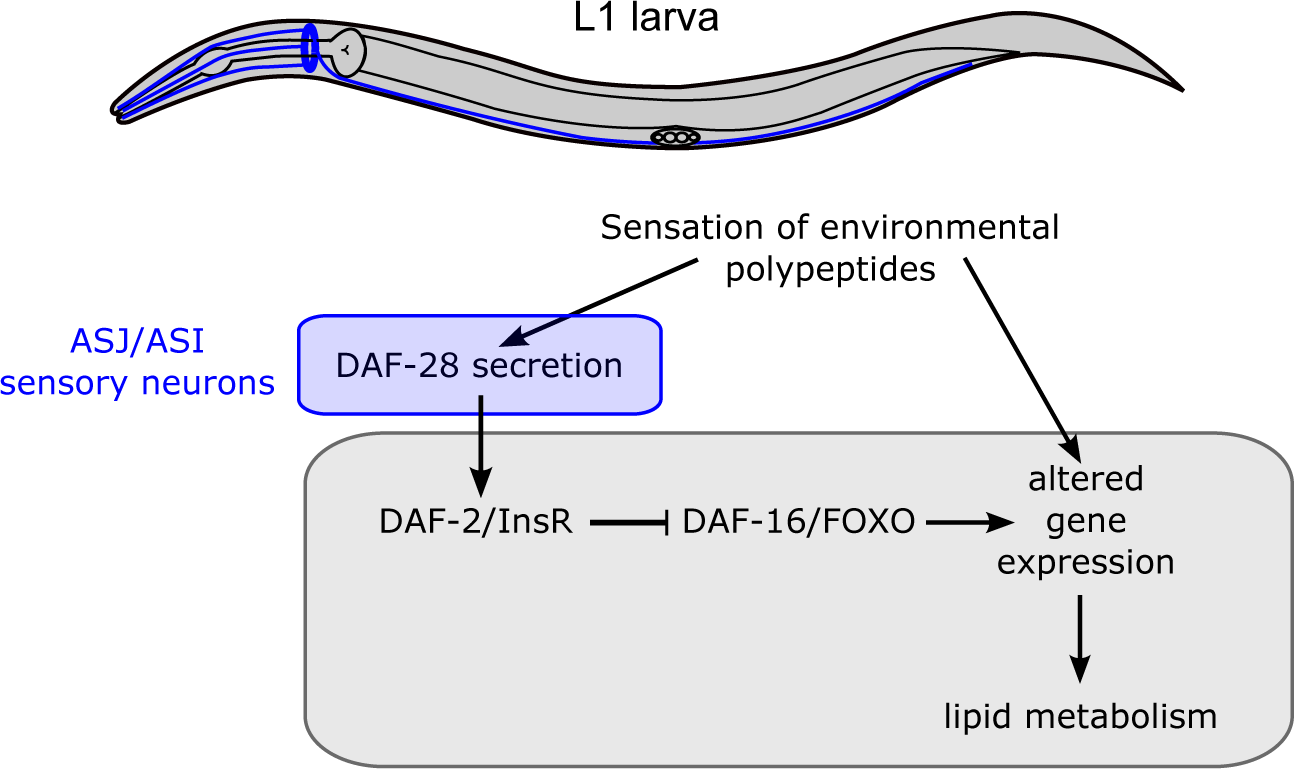
An organismal response to perception of food. Environmental polypeptides sensed by chemosensation activate IIS and drive altered gene expression, which affects lipid metabolism.

Since the relevant modality of perception appeared to be chemosensory, we wanted to identify a molecular component of bacterial food that functions as an environmental cue for the worm. We found that LB medium, a common nutrient broth for culturing *E. coli*, as well as its components, yeast extract and tryptone, caused irreversible arrest in worms exposed to them in the presence of ivermectin, similar to the effect of HB101 (Fig. 5B). Yeast extract results from autolysis of *S. cerevisiae* and contains a complicated mixture of amino acids, peptides, carbohydrates, and vitamins. Tryptone is a tryptic digest of the protein casein, resulting in polypeptides of varying lengths. Since tryptone is much simpler than yeast extract, we decided to focus our investigation there. We tested undigested casein and casamino acids, which is casein that has been through acid hydrolysis to produce free amino acids. We also tested bovine serum albumin (BSA) as another form of protein. We found that casein and BSA significantly prevented recovery while casamino acids did not (Fig. 5B). Since casamino acids do not contain polypeptide, these results suggest polypeptide is perceived. We also tested a solution of the ten essential amino acids for *C. elegans*, ethanol, glucose, and a combination of all three, and found that none of these significantly affected recovery (Supp. Fig. 5A). Perception of polypeptides and other potential food cues also caused GFP::DAF-16 to translocate to the cytoplasm (Supp, Fig. 5B,C). When otherwise starved larvae were permitted to ingest polypeptide or other potential cues, they supported survival (Supp. Fig. 5D) but not development (based on the M-cell division assay; data not shown), as if providing an incomplete source of nutrition. This treatment also compromised the ability of larvae to subsequently recover in standard culture conditions (Supp. Fig. 5E,F), reminiscent of the effect of exposure to food in the presence of ivermectin. In summary, we conclude that starved worms perceive polypeptides, as if they are a food-associated cue, though other cues may also be involved.

We wanted an alternative and more ecologically relevant approach than using ivermectin to determine if starved worms perceive polypeptide. Dauer larvae have an internal plug blocking the pharynx and do not pump (Cassada and Russell 1975, Riddle, Swanson et al. 1981). Tryptone and LB promoted dauer recovery, as did HB101, while casamino acids and the buffer S-complete did not (Fig. 5C). These results further support the conclusion that *C. elegans* perceive environmental polypeptides when starved as a food-associated cue, and they suggest that such perception provides an important regulatory input for dauer recovery.

## Discussion

We sought to uncouple the effects of food perception and ingestion on *C. elegans* development, gene expression and metabolism. We report that perception is not sufficient to promote development, but that it activates IIS and alters gene expression and lipid metabolism. We also report that starved larvae sense environmental polypeptides, as if worms use them as a food-associated cue to anticipate feeding and development.

The most striking phenotype we report is the irreversible developmental arrest of larvae that are starved in the presence of food, so that they perceive food without eating it. Ivermectin binding has been characterized as irreversible (Cully, Vassilatis et al. 1994, Vassilatis, Arena et al. 1997, Horoszok, Raymond et al. 2001). These studies involved very different time scales from ours, and they used ivermectin doses 50-100-fold greater than us. Nonetheless, we considered irreversible binding as an explanation for irreversible arrest, but several lines of evidence suggest otherwise. Worms exposed to the relatively low dose of ivermectin we used without food almost completely recover. Also, recovery was rescued by blocking transcription or translation. In addition, we see a similar reduction in recovery rate in otherwise starved L1 larvae exposed to food cues. This observation along with the effect of ivermectin and food suggest that perception of food cues without ingestion of complete nutrition underlies the irreversible arrest phenotype. We speculate that perception of food alters metabolism to prime the animal for feeding and development, but that the changes that occur are detrimental if not accompanied by feeding.

We present evidence that food perception elicits a gene expression response that is largely subsumed by the feeding response, and that this response is in part due to activation of IIS. Notably, the gene expression response and activation of IIS were relatively transient, as if larvae initially respond to food perception but this response is not maintained without feeding and ingestion of nutrients. We imagine that the transient nature of this response is due to habituation of perception or antagonism from internal starvation signals, or a combination of the two. Up-regulation of IIS during L1 starvation promotes cell division (Chen and Baugh 2014), but perception of food did not, though IIS was activated. We believe the transient nature of IIS activation by food perception explains the lack of postembryonic development. Despite the transient nature of the responses to food perception, they nonetheless have physiological consequences as demonstrated by accumulation of lipid droplets and irreversibility of developmental arrest.

Insulin-like peptides *daf-28, ins-6* and *ins-4* govern postembryonic development, and their transcription is positively regulated by nutrient availability (Li, Kennedy et al. 2003, Cornils, Gloeck et al. 2011, Chen and Baugh 2014). We show that *daf-28* transcription is up-regulated by perception of food, and that it plays a specific role in activating IIS in response to perception. That is, *daf-28* was specifically required for food perception to cause GFP::DAF-16 translocation to the cytoplasm, though it was dispensable for translocation in response to feeding, suggesting overlapping function with other insulin-like peptides. We identify polypeptides as a bacterial component that functions as a food cue for starved larvae. Perception of polypeptide caused GFP::DAF-16 to translocate and caused an irreversible arrest phenotype. Together our results suggest that chemosensation of environmental polypeptides promotes transcription and likely secretion of DAF-28 from ASI and ASJ amphid neurons to mediate systemic effects on gene expression and metabolism. In support of a direct effect of food perception on insulin-like peptide secretion from chemosensory neurons, inhibiting translation with cycloheximide did not block the effect of perception on GFP::DAF-16 localization. However, activation of IIS did not account for the entire gene expression response to food perception, nor did it account for the irreversible arrest phenotype. We conclude that food perception affects additional signaling pathways, and that these pathways collaborate with IIS to regulate gene expression and metabolism.

We conclude that *C. elegans* larvae sense environmental food-associated cues such as polypeptides, and that this perception affects signaling, gene regulation and metabolism. Worms likely use chemosensation to find food, and food perception may also serve to prime starved larvae for feeding and development. Such priming is apparently detrimental if not accompanied by feeding within hours, but we believe such a scenario where food cues are present without food is unnatural. In contrast, dauer larvae represent a common situation where starved larvae rely on perception to regulate development and metabolism. We show that dauer larvae exit arrest and resume development in response to perception of environmental polypeptides, similar to their response to NAD^+^ (Mylenko, Boland et al. 2016). Starved non-dauer larvae are able to feed immediately upon encountering food, but perception of environmental cues could accelerate the organismal response by not requiring ingestion and assimilation of nutrients. With a fluctuating food supply and boom and bust population dynamics, we believe metabolic priming via food perception contributes to fitness by accelerating recovery from developmental arrest.

## Materials and Methods

### *C. elegans* growth conditions and strains

Strains were maintained on agar plates containing standard nematode growth media (NGM) seeded with *E. coli* OP50 at 20°C. The wild-type strain N2 (Bristol) and the following mutants and transgenes were used: *daf-2(e1370), daf-16(mu86), ayIs7[Phlh-8::GFP], avr-14(vu47), glc-3(ok321), avr-15(vu227), glc-1(pk54), dukIs9[Pmyo-2::avr-15*+*Pmyo-2::mCherry*+*Pajm-1::AJM-1::GFP], dukIs10[Pmyo-2::avr-15*+*Pmyo-2::mCherry*+*Phlh-8::GFP], qyIs288 [Pdaf-16::GFP::DAF-16* + *unc-119(*+*)], qyIs289 [Pdaf-16::GFP::DAF-16* + *unc-119(*+*)], daf-28(tm2308), ins-4, 5, 6(hpDf761)*. Standard genetic techniques were used to make combinations of alleles.

*dukIs9* injection mix contained the following: 1 ng/µL pCFJ90 (*Pmyo-2::mCherry*), 1 ng/µL pPD30_69_TK414_4A (*Pmyo-2::avr-15*), and 50 ng/µL pJS191 (*Pajm-1::AJM-1::GFP*). *dukIs10* injection mix contained the following: 1 ng/µL pCFJ90 (*Pmyo-2::mCherry*), 1 ng/µL pPD30_69_TK414_4A (*Pmyo-2::avr-15*), and 50 ng/µL pJKL464 (*Phlh-8::GFP*).

### Hypochlorite treatment and L1 arrest assays

Mixed-stage cultures on 10 cm NGM plates were washed from the plates using virgin S-basal (S-basal lacking ethanol and cholesterol) and centrifuged. A hypochlorite solution (7:2:1 ddH_2_O, sodium hypochlorite (Sigma), 5 M KOH) was added to dissolve the animals. Worms were centrifuged after 1.5-2 minutes in the hypochlorite solution and fresh solution was added. Total time in the hypochlorite solution was 8-10 minutes. Embryos were washed three times in virgin S-basal buffer (no ethanol or cholesterol) before final suspension in 3 to 6 mL virgin S-basal at a density of 1 worm/µL. Ivermectin (Sigma) dissolved in DMSO was added to the appropriate cultures. Ivermectin dose was adjusted such that worms did not eat. The dosage was 10 ng/mL unless otherwise stated. *avr-14(vu47); glc-3(ok321) avr-15(vu227) glc-1(pk54); dukIs10[Pmyo-2::avr-15*+*Pmyo-2::mCherry*+*Phlh-8::GFP]* was treated with 20 ng/mL ivermectin, or 22.85 nM. *avr-14(vu47); glc-3(ok321) avr-15(vu227) glc-1(pk54); dukIs9[Pmyo-2::avr-15*+*Pmyo-2::mCherry*+*Pajm-1::AJM-1::GFP]* was treated with 50 ng/mL ivermectin. Different doses of ivermectin were used to adjust for different levels of ivermectin resistance in different strains. DMSO was added in equal amounts to control tubes. DMSO concentration ranged from 0.05% to 0.2%. Embryos were cultured in a 16 mm glass tube on a tissue culture roller drum at approximately 25 rpm and 21-22°C.

For the M-cell division assay, 1 day following the hypochlorite treatment above the worms were put in the appropriate condition (LB, tryptone, etc.) and cultured for 7 days before 100 larvae per replicate were examined on a slide on a compound fluorescent microscope. For the seam cell division assay, 1 day following the hypochlorite treatment above HB101 was added at 25 mg/mL for 2 days and the V1-6 cells on one side of the animal were scored for 60 larvae per replicate.

### Starvation recovery

Animals were treated in hypochlorite solution and suspended in virgin S-basal with DMSO or ivermectin as described above. One day after hypochlorite treatment, the appropriate bacteria (HB101 unless otherwise stated) or partial food was added at the appropriate dose (25 mg/mL for bacteria unless otherwise stated). HB101 filtrate was created by filtering HB101 at 25 mg/mL through a 22 µm filter. Yeast extract was at 5 mg/mL. Tryptone and casamino acids were at 10 mg/mL. Due to solubility limitations, casein and BSA were at 1 mg/mL. Ethanol was at 0.095% (v/v). Glucose was at 5% (w/v), or 278 mM. Amino acid solution (16 mg/mL) made up as in (Fukuyama, Kontani et al. 2015). Food addition was considered the 0 hr timepoint (Fig. 1A). 100 µL aliquots were sampled at the stated times up to 24 hr and placed around the edge of a HB101 lawn on NGM plates. Number of plated worms (T_p_) was counted and the plates were incubated at 20°C. After three days the number of animals that recovered to at least the L4 stage (T_R_) was counted. Recovery was calculated as T_R_/T_p_.

### GFP bead ingestion

Cultures were setup as for a starvation recovery experiment as above, except instead of plating the worms after 24 hr GFP beads (Fluoresbrite^®^ YG Carboxylate Microspheres 0.10µm from Polysciences) were added at 1:200 to the cultures. After 3-4 hr the cultures were examined on a slide on a compound fluorescent microscope. The location of the GFP beads was scored for 40 worms per replicate.

### α-amanitin and cycloheximide treatment

Dose response curves with α-amanitin (Sigma) and cycloheximide (Sigma) were done using the gpIs1 [*Phsp-16.2*::GFP] reporter (Link, Cypser et al. 1999) to find a dose that prevented fluorescence in response to heat shock at 33°C for two hr. These doses were determined to be 5 mM for cycloheximide and 25 µg/mL for α-amanitin. Both drug stocks were dissolved in water. The starvation recovery assay was set up as above, with drugs added two hr before food addition and cultures washed three times with 10 mL virgin S-basal before plating.

### mRNA-Seq and associated analysis

Worm cultures for *avr-14(vu47); glc-3(ok321) avr-15(vu227) glc-1(pk54); dukIs10[Pmyo-2::avr-15*+*Pmyo-2::mCherry*+*Phlh-8::GFP]* were set up using the hypochlorite treatment as described above, except in S-complete and scaled up to 20 mL per condition. Either ivermectin was added at 5 ng/mL or DMSO was added at 0.1%. After 24 hr to allow for hatching and synchronization, HB101 was added at 25 mg/mL to the food tubes. Samples were collected at 1 hr and 24 hr after food addition. To collect the samples, worms were washed 3 times with 10 mL virgin S-basal then concentrated in 100 uL and frozen in liquid nitrogen. RNA was extracted with Trizol and chloroform. Libraries were prepared for sequencing using the NEBNext Ultra RNA Library Prep Kit for Illumina (E7530) with 250-400ng of starting RNA per library and 13 cycles of PCR. Libraries were sequenced using Illumina HiSeq 4000. Bowtie was used to map reads to the WS210 genome (Langmead, Trapnell et al. 2009). Transcripts annotated in WS220 that were mapped to the WS210 genome coordinates were also included, as described previously (Maxwell, Antoshechkin et al. 2012). Mapping efficiencies ranged from 78-85% for all libraries. HTSeq was used to generate count tables for each library (Anders, Pyl et al. 2015). Count tables were analyzed for differential expression using the edgeR package in R (Robinson, McCarthy et al. 2010). Detected genes were considered those expressed at a level of at least 1 count-per-million (CPM) in at least four libraries, reducing the number of genes included in the analysis to 18190. The “calcNormFactors” function was used to normalize for RNA composition and the tagwise dispersion estimate was used for differential expression analysis. The exact test was used for pairwise comparisons of conditions. Differentially expressed genes were considered those with an FDR < 0.05 and with |log_2_ (fold change)| > 0.5. Principal component analysis was performed using all libraries and all genes used in differential expression analysis (18190 genes). Counts-per-million (CPM) values for each gene were mean-normalized across all libraries and log_2_ transformed prior to using the prcomp function in R. GEO accession number for the dataset is GSE114955. GO term analysis was performed using GOrilla (Eden, Lipson et al. 2007, Eden, Navon et al. 2009). AmiGO 2 was accessed to download the genes in the metabolic process GO term (GO:0044710) and the larval development GO term (GO:0002164) (Ashburner, Ball et al. 2000, Carbon, Ireland et al. 2009, The Gene Ontology 2017).

### GFP:: DAF-16 localization

The qyIs288 *[Pdaf-16::GFP::DAF-16* + *unc-119(*+)] and qyIs289 *[Pdaf-16::GFP::DAF-16* + *unc-119(*+)] reporters (Kaplan, Chen et al. 2015) were analyzed in a *daf-16(mu86); unc-119(ed4)* mutant background. Standard genetic methods were used to cross *daf-28(tm2308)* and *ins-4, 5, 6(hpDf761)* into this background as well. Cultures were set up using the hypochlorite treatment as described above. One day later HB101 was added at 25 mg/mL. One hr or 24 hr after food addition 50 larvae per replicate were examined on a slide on a compound fluorescent microscope.

### Reporter gene analysis

The mgIs40 [P*daf-28*::GFP] reporter (Li, Kennedy et al. 2003) was analyzed in a wild type genetic background. Strain was maintained on NGM agar plates with *E. coli* OP50 as food at 20°C. Eggs were prepared by standard hypochlorite treatment. These eggs were used to set up a liquid culture consisting of virgin S-basal with a defined density of 1 worm/µl. Ivermectin was added at 10 ng/mL to the appropriate cultures. After 18 hr to allow for hatching, the *E. coli* HB101 was added at 25 mg/ml to the fed samples. 6 hr post food addition, the samples were washed three times with 10 mL S-basal and then run through the COPAS BioSorter measuring GFP fluorescence. Analysis of the COPAS data was performed in R. Data points were removed if they were determined to be debris by size. Fluorescence signal was normalized by optical extinction. For imaging, the samples were prepared in the same way then paralyzed with 3.75 mM sodium azide and placed on a Noble agar slide. Images were taken on a compound fluorescent microscope.

### Fixation and Nile red staining

Cultures of N2 wild type were setup as for a starvation recovery experiment as above, except instead of plating the worms after 24 hr the cultures were washed three times with 10 mL virgin S-basal. Worms were concentrated in approximately 100 µL and frozen at −80°C. Fixation and staining protocol was modified from (Pino, Webster et al. 2013), using 1.7 mL Eppendorf tubes instead of 96-well plates and 200 µL solution additions instead of 150 µL. Images were taken on a compound fluorescent microscope. Fat droplets were quantified using the Analyze Particles function in Image J. Images were thresholded using negative controls to remove background. Minimum particle size was set as 0.002 in^2.^

### Starvation survival

N2 wild type animals were treated in hypochlorite solution and suspended in virgin S-basal or the appropriate media as described above. 100 µL aliquots were sampled on different days and placed around the edge of an OP50 lawn on NGM plates. Number of plated worms (T_p_) was counted and the plates were incubated at 20°C. After two days the number of animals that survived (T_s_) was counted. Survival was calculated as T_s_/T_p_. Survival curves were obtained by fitting survival data for each trial with the function

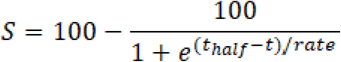

### Quantitative image analysis of size

N2 wild type animals were treated in hypochlorite solution and suspended in virgin S-basal or the appropriate media as described above. At the 50% survival times determined from starvation survival experiments the worms were spun down at 3000 rpm for 1 minute and pellets were transferred to OP50 seeded NGM plates. Worms were allowed to recover for 48 hr at 20°C. Worms were then imaged and images were processed using the WormSizer plug-in for Fiji/ImageJ as described (Moore, Jordan et al. 2013).

### Dauer recovery

N2 worms were treated in hypochlorite solution as described above then resuspended in S-complete at a concentration of 5 worms/µL and 1 mg/mL HB101 in 25 mL Erlenmeyer flasks (Baugh, Kurhanewicz et al. 2011). Flasks were placed on a shaker at 20°C for one week to form dauers. Cultures were spun down at 3000 rpm for 1 minute. Supernatant was aspirated and the appropriate media (LB, tryptone, etc.) was added, retaining concentration of 5 worms/µL. Cultures were returned to shaker for three days. Approximately 75-100 worms were placed on a depression slide and scored as dauer, L4, or adult.

### Data analysis and statistics

Data were handled in R and Excel. Graphs were plotted in the R packages ggplot2 or Vennerable or Excel. Statistical tests were performed in R or Excel. Starvation survival analysis was performed on 50% survival times (*t_half_*), which were obtained as in (Kaplan, Chen et al. 2015), with unpaired t-tests performed where n = number of replicates.

## Acknowledgements

We would like to thank Joel Meyer and David R. Sherwood for sharing lab equipment and Baugh lab members for helpful discussions. We thank the Duke University School of Medicine and the Center for Genomic and Computational Biology for use of the Sequencing and Genomic Technologies core resource, which provided RNA sequencing service. Some strains were provided by the CGC, which is funded by NIH Office of Research Infrastructure Programs (P40 OD010440). Funding was provided by the National Institutes of Health (R01GM117408, LRB).

## Author Contributions

Contributed reagents: REWK, JAD. Conceived and designed experiments: REWK, JAD, LRB. Performed experiments: REWK, RC. Analyzed data: REWK, AKW. Wrote the paper: REWK, LRB.

## Figure Legends

**Supp. Figure 1.**
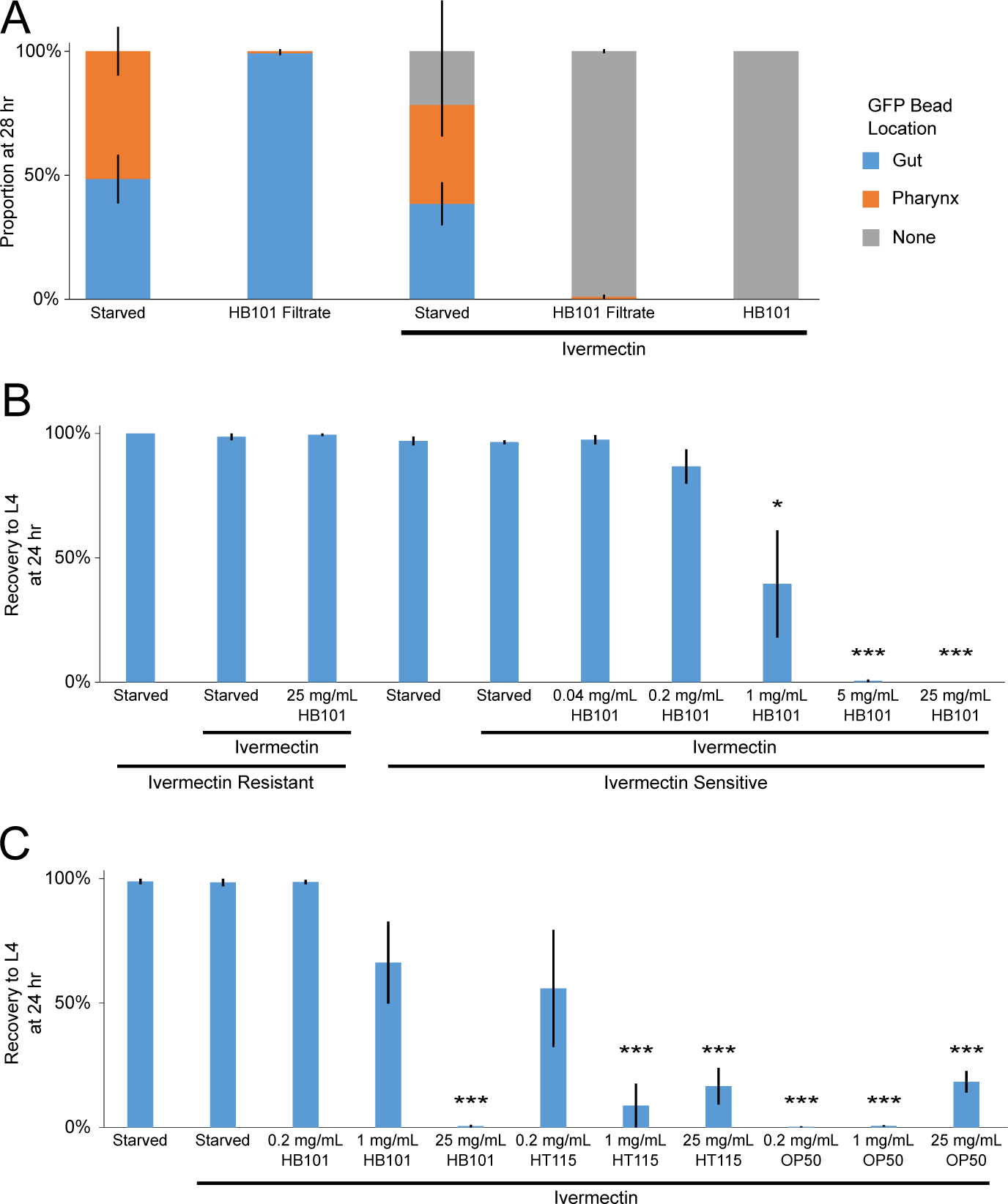
Further characterization of ivermectin system and starvation recovery. (A) The proportion of larvae that displayed the stated localization of GFP beads three to four hours after bead addition is plotted for three biological replicates. (B-C) The proportion of larvae that recovered to at least the L4 stage after three days of recovery is plotted for three to four biological replicates. Ivermectin Resistant = *avr-14(vu47); glc-3(ok321) avr-15(vu227) glc-1(pk54)*. Ivermectin Sensitive = *avr-14(vu47); glc-3(ok321) avr-15(vu227) glc-1(pk54); dukIs10[Pmyo-2::avr-15*+*Pmyo-2::mCherry*+*Phlh-8::GFP]*. ***p<0.001, *p<0.05; unpaired t-test. (A-C) Error bars are SEM.

**Supp. Figure 2.**
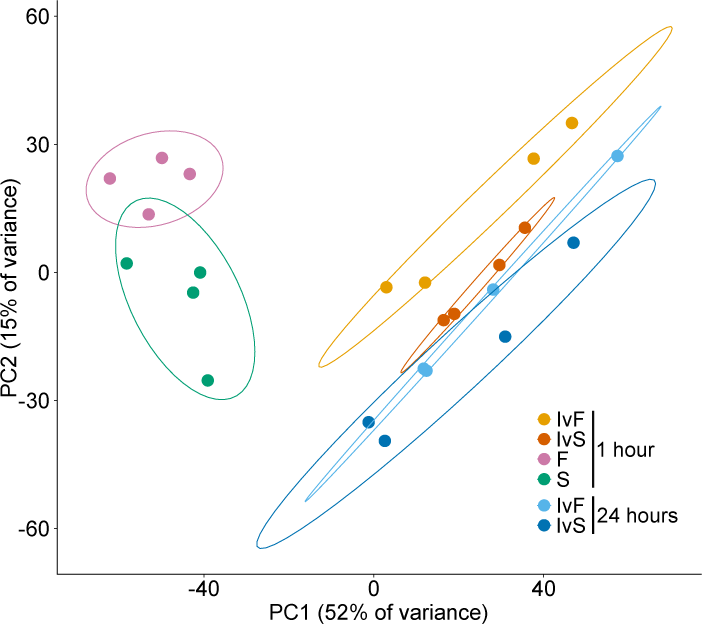
Ivermectin affects transcription in larvae. PCA of four biological replicates is plotted. Ellipses represent 80% confidence interval.

**Supp. Figure 3.**
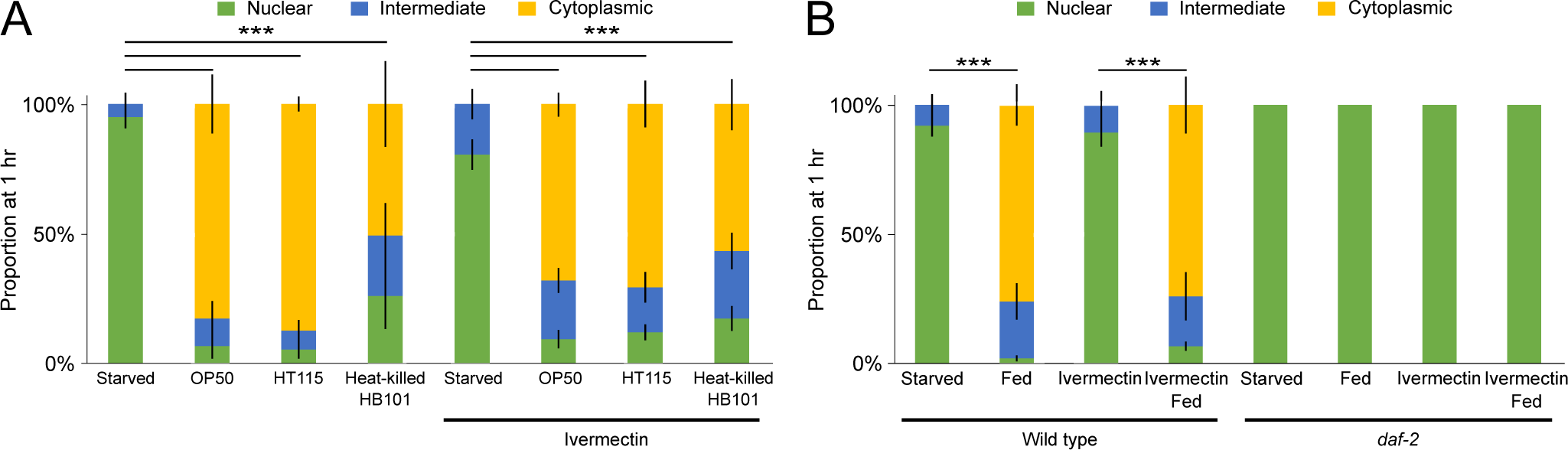
GFP::DAF-16 localization responds to perception of many bacterial foods and requires *daf-2*. (A-B) GFP::DAF-16 localization is plotted for three to four biological replicates. Wild type is in N2 background. ***p<0.001; unpaired t-test. Error bars are SEM.

**Supp. Figure 4.**
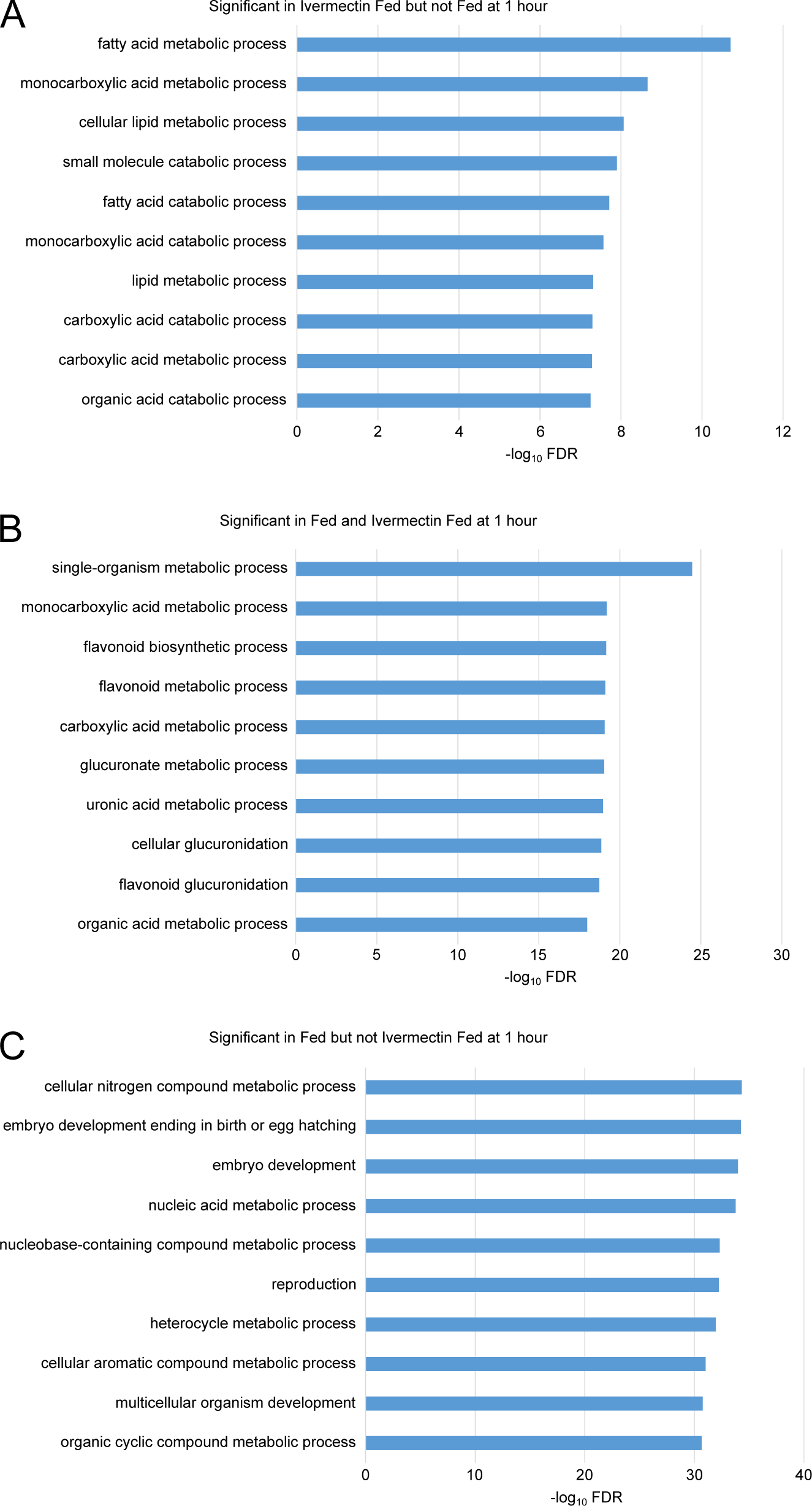
Food perception affects metabolism-related GO terms. (A-C) Top ten GO terms for stated gene groups are plotted from four biological replicates of RNA-seq.

**Supp. Figure 5.**
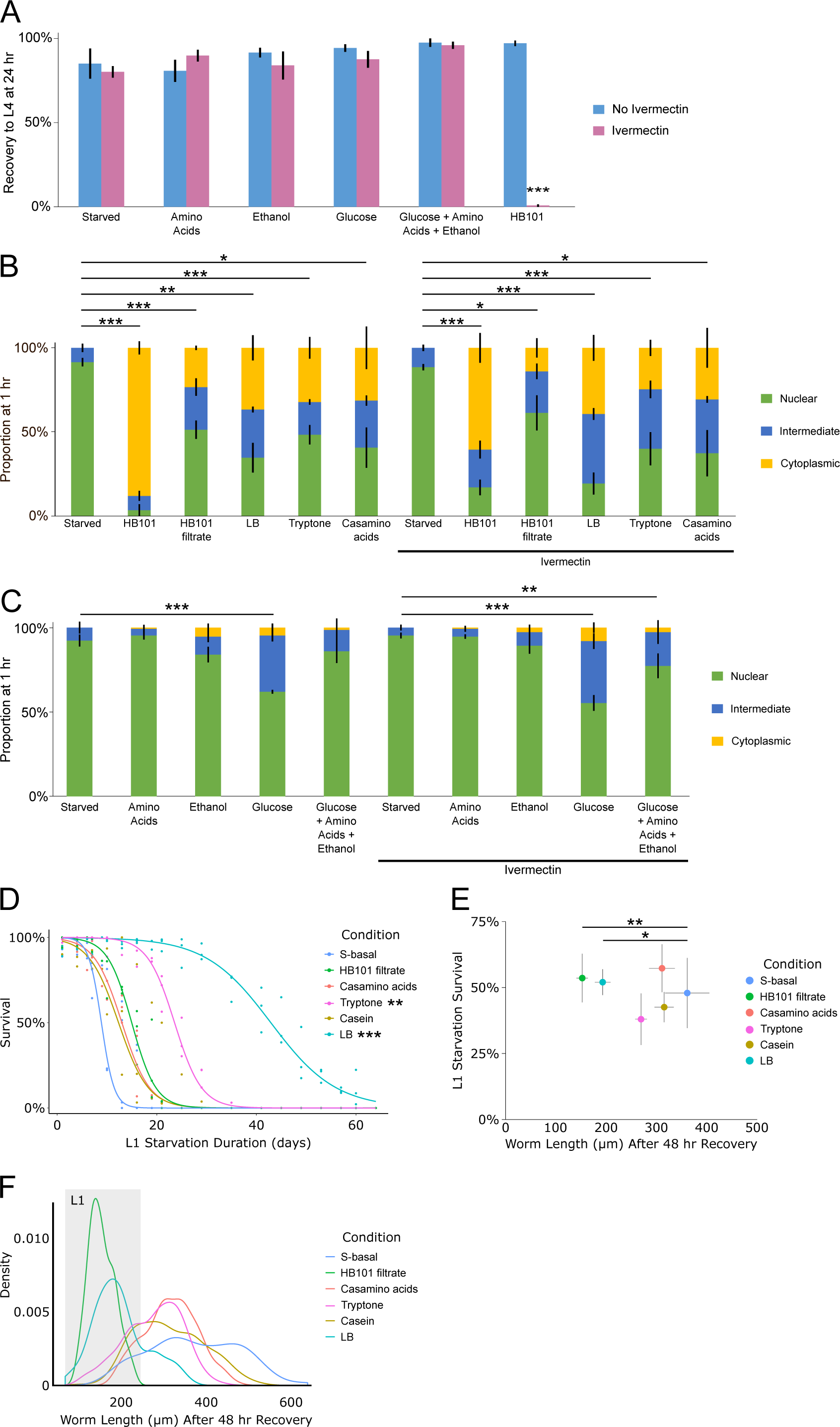
Perception and physiological effects of potential food cues. (A) The proportion of larvae that recovered to at least the L4 stage after three days of recovery is plotted for three biological replicates. (B-C) GFP::DAF-16 localization is plotted for three to six biological replicates. (D) L1 starvation survival is plotted over time. A logistic regression of mean survival from three biological replicates is shown. (E) Worm length following 48 hr of recovery is plotted relative to L1 starvation survival. (F) Worm length following 48 hr of recovery is plotted as a density plot, showing altered population composition. (A-E) ***p<0.001, **p<0.01, *p<0.05; unpaired t-test. Error bars are SEM.

S1 Dataset. **mRNA-seq analysis of food perception.** Results of mRNA-seq analysis, including counts per gene, logFC, FDR, GOrilla analysis, and GO term gene lists used are included.

